# Shared and distinct microRNA profiles between HT22, N2A and SH-SY5Y cell lines and primary mouse hippocampal neurons

**DOI:** 10.1101/2025.06.01.657291

**Authors:** Ronan Murphy, Javier Villegas-Salmeron, Amaya Sanz-Rodriguez, Tobias Engel, Catherine Mooney, David C Henshall, Eva M Jimenez-Mateos

**Affiliations:** Discipline of Physiology. School of Medicine. Trinity College Dublin. The University of Dublin, Dublin, Ireland; Department of Physiology and Medical Physics, RCSI University of Medicine and Health Sciences, Dublin, Ireland; FutureNeuro Research Ireland Centre, RCSI University of Medicine and Health Sciences, Dublin, Ireland; School of Computer Science, University College Dublin, Dublin, Ireland

**Keywords:** Brain disease, Epilepsy, immortalized cell lines, microRNAs, neurons, small noncoding RNA

## Abstract

MicroRNAs (miRNA) are small non-coding RNAs that are key negative regulators of gene expression. Their roles include shaping the gene expression landscape during and after brain development by defining and maintaining levels of proteins that generate the distinct morphological and functional properties of neurons and other brain cell types. HT22, N2A, and SH-SY5Y are common immortalized neuronal cell lines that offer simple, less expensive, and time-saving options for *in vitro* modelling to evaluate miRNA functions. The extent to which these lines reflect primary neurons remains, however, unclear. Here, we benchmarked the miRNA profiles of cultured mouse hippocampal neurons against Argonaute-loaded miRNAs in the adult mouse hippocampus and miRNA data from the hippocampus of patients with drug-resistant temporal lobe epilepsy. We then compared the miRNA expression landscape in HT22, N2A and SH-SY5Y against mouse hippocampal primary cell cultures. We profiled over 700 miRNAs across the lines and detected 310 miRNAs in all four cell types. This included detection of neuron-enriched miRNAs such as miR-124 and miR-128, although the cell lines typically displayed lower levels of these than in primary neurons and reference adult hippocampal tissue. The miRNA profile in the HT22 cell line showed the highest correlation to the mouse primary neuronal cultures. Together, this study provides a dataset on basal miRNA expression across commonly used cell lines for neuroscience research and evidence for both conserved and distinct profiles that should be used to inform decisions on cell lines for modelling brain and miRNA research.

## Introduction

MicroRNAs (miRNA) are small non-coding RNAs (22-24 nucleotides) that post-transcriptionally regulate gene expression. This is achieved by the miRNA being loaded into the binding pocket of an Argonuate (Ago) protein, thereby forming an RNA-induced silencing complex (RISC) that recognizes complementary sequences in the 3’ untranslated region of the targeted messenger RNA (mRNA) [1]. MiRNAs are master regulators of gene expression, with individual miRNAs able to regulate the translation of up to hundreds of protein-coding mRNAs. The impact of a single miRNA on the protein level of a target is, however, generally modest, suggesting that cells use miRNAs to fine-tune their protein landscape [2]. MiRNAs are essential for organism development and are key regulators of cellular identity, with disruption of miRNA production resulting in embryonic lethality in worms, flies, zebrafish, and mice, mainly due to neuronal defects [2, 3]. In the adult brain, miRNAs remain essential for homeostasis, with disruption of miRNA production leading to neurological and neurodegenerative conditions [4, 5], including epilepsy [6]. In the brain, miRNAs show distinct expression patterns specific to developmental stages, brain regions, and cell types [7–10], reflecting their critical role in neuronal differentiation and establishing and maintaining neuronal diversity [11, 12]. For example, miR-124 is key for establishing the neuronal lineage during embryogenesis via regulation of the transcription factor REST and suppressing non-neuronal transcripts in neurons [13–15]. MiR-125 is critical for neuronal maturation and maintaining neuronal properties [16], and miR-133b is expressed in and regulates the maturation of dopaminergic neurons in the midbrain [17]. Among key processes shaped by the actions of miRNAs are neuronal structure and function, including dendritic morphology and the complement of neurotransmitter receptors, ion channels, and transporters [2]. Consequently, miRNAs have been extensively studied as potential mediators of aberrant gene expression in neurological diseases [2, 18].

Immortalized cell lines are commonly used to understand and identify molecular and cellular processes. In classical studies, HeLa cells were used to demonstrate that miR-124 is critical to establishing the neuronal gene expression landscape [19]. Cell lines are an invaluable tool that can provide a simple way to identify and test molecular and cellular mechanisms before moving to primary cultures or *in vivo* experiments [20–23]. However, immortalized cell lines typically have a tumoral origin or have been modified to be kept cultured indefinitely, which results in phenotypes and patterns of gene regulation that do not reflect some neuronal characteristics and responses to stimuli. Examples of common immortalized neuronal cell lines are the HT22, N2A and SH-SY5Y cells. HT22 cells are immortalized primary mouse hippocampal neuronal cells, derived from parent HT4 cells [22, 24]. Undifferentiated HT22 cells express low levels of cholinergic markers and glutamate receptors [25, 26], in contrast to mature hippocampal neurons. Differentiated HT22 cells express higher levels of *N*-methyl-D-aspartate receptor (NMDAR) mRNA [27], making them susceptible to glutamate-induced excitotoxicity [26]. N2A is a mouse neuroblastoma cell line, commonly used to study neuronal differentiation and used extensively to test novel cancer treatments [28]. SH-SY5Y are neuroblastoma cells of human origin with a neuroblast-like morphology and expression of immature neuronal markers [29]. After differentiation with retinoic acid and neurotrophins, SH-SY5Y cells become morphologically similar to primary neurons and express neuron-specific markers [29].

Previous studies have evaluated the expression of miRNAs in cortical primary neurons and glial cells (astrocytes, microglia and oligodendrocytes) [8]. This identified a set of miRNAs that are enriched in neurons, with expression fivefold higher than the glial cells, including miR-124, miR-132 and miR-135b, while the same cells display very low expression of miR-21 and miR-146a compared to glial cells [8]. Importantly, neurological disorders can result in altered miRNA expression, contributing to pathology [30]. Here, cell lines have contributed to discoveries on miRNAs, including linking miR-134 to the pathophysiology of epilepsy [31–37].

While the miRNA profiles of neuronal cell lines have been described, researchers typically report on individual cell lines. Having a catalogue of miRNA profiles of multiple cell lines analysed together is important for the selection of the most appropriate cell line for a particular research project. Here, we established the basal miRNA profile for three commonly used immortalized cell lines in neuroscience research and compared these to the profile in primary hippocampal neurons from mice and brain data. The data will support decision-making on the best cell line to study specific miRNAs or molecular mechanisms before moving to *in vivo* experiments.

## Materials and Methods

### Primary cell culture

Primary cultures of hippocampal neurons were prepared following a previously described protocol [34]. E18 embryonic mouse brains were removed and placed into a solution of HBSS (Sigma-Aldrich Ireland Ltd., Wicklow, Ireland). Hippocampi were dissected and cells were disaggregated using papain (100 units/ml, Sigma-Aldrich Ltd., Wicklow, Ireland) for 30 min at 37 °C. Neurons were then isolated by tissue dissociation and plated onto a 1 mg/ml poly-L-lysine and 20 μg/ml laminin. Cells were maintained in Neurobasal medium supplemented with B-27 and N2 (Thermo-Fisher, MA, USA) at 37 °C in a humidified atmosphere with 5% (v/v) CO2 for 10 days. Half of the medium was changed to fresh medium every second day. RNA was isolated at day *in vitro* 10 (DIV10) as described below/

### Cultured cell lines

HT22 cells (a gift from Prof C. Culmsee) were grown in Dulbecco’s Modified Eagle Medium (DMEM, Lonza Biologics, UK), 4.5 g/L high glucose with glutamine and sodium pyruvate containing 10% FBS (fetal bovine serum). SH-SY5Y cells (a gift from Prof J.H.M Prehn) were cultured in Dulbecco’s Modified Eagle’s Medium-F12. N2A cells (a gift from Prof J.H.M Prehn) were cultured in a supplemented DMEM medium (ThermoFisher, MA, USA). All immortalized cell lines were supplemented with penicillin/streptomycin, and incubated in 5% CO_2_ humidified atmosphere at 37°C. The immortalized cell lines were plated at a 10.000 cells/mm^2^ density in an M24 plate and harvested 48 hours later. For triplicates, the vial was defrosted in three independent plates and maintained as three independent cell cultures. After four passages, cell lines were seeded in an M24 plate, and RNA was extracted as described below.

### RNA extraction and OpenArray

Total mRNA was extracted using the Trizol protocol as previously described [34, 38]. Quantity of mRNA was measured using a Nanodrop Spectrophotometer (Thermo Fisher Scientific, Rockford, IL, USA), and RNA dilutions were made up in nuclease-free water. 100ng of purified RNA was processed by reverse transcriptase and pre-amplification steps following the manufacturer’s protocol (Applied Biosystems). The pre-amplification reaction was mixed with TaqMan OpenArray Real-Time PCR Master mix (1:1). The mix was loaded onto the OpenArray rodent microRNA panel (750 mouse/rat miRNAs (Cat number: 4461105)) and ran using a QuantStudio 12K Flex PCR (Life Technologies).

### Analysis of OpenArray

Analysis and statistical analysis of the OpenArray miRNA profiling data were performed in R. Data were first filtered according to amplification score (AmpScore >= 1.24), quantification cycle confidence (Cqconf >= 0.8) and cycle threshold (Ct > 10 and < 35). For the comparison of the expression across the four cell types, miRNAs were included if they were present in at least 2 of the 3 samples in each group. Missing data points were imputed as the mean Ct of the other two samples, and the data were quantile normalised using the Bioconductor package qpcrNorm [39]. Principal component analysis (PCA) was performed on the quantile normalised data. Heatmaps and Venn diagrams were generated using the Complex Heatmap [40] and Eulerr [41] packages in R, respectively. Correlation was calculated using the Pearson correlation coefficient. Correlation graphs, slope graphs and principal component analysis (PCA) plots were generated using ggplot2 [42]. Differential expression analysis was performed using the limma package [43]. p-values were adjusted for multiple testing by controlling the false discovery rate (FDR) according to the method of Benjamini and Hochberg [44]. A miRNA was considered to be differentially expressed if the adjusted p-value was < 0.05. Relative expression was analysed using the ΔΔCt method with all miRNA analysis normalised by subtracting the geometric mean Ct for each miRNA. Statistical analysis on individual miRNAs was performed using ANOVA followed by Dunnet test. Significance was assigned as p < 0.05.

### RNA-Seq data analysis

Analysis of RNA-seq data was performed using the Bioconductor package edgeR [45]. Data were filtered by miRNA counts per sample (count > 10), and only samples with ∼ 8 million reads were used for the analysis. Only miRNAs present in all 3 samples were included in the analysis. Data were standardised by log counts per million followed by a linear transformation to convert the count data to the equivalent Ct range (10–35). Next, the data were merged with the qPCR data and quantile normalised using the Bioconductor package qpcrNormand [39]. Differential expression analysis was performed as described above.

## Results

### Cultured mouse primary hippocampal neurons have a similar miRNA profile to Ago2-bound miRNAs from the adult mouse hippocampus

Mouse primary hippocampal neurons were selected as a suitable comparator for the miRNA profiles of HT22, N2A and SH-SY5Y cells (Table 1). First, and using previously published data [38, 46], we compared our in vitro hippocampal primary neurons from mice (DIV (Day in vitro) 10) to the miRNA profile of the mouse hippocampus by comparing with Ago-2 bound miRNAs from the hippocampus of adult C57BL/6 mice [38]. We also compared the profile to miRNAs in the adult human hippocampus [46]. When compared with mouse tissue, we found that 78% of the expressed miRNAs in our primary mouse neurons were also among those actively loaded in the RISC in the mouse brain (Figure 1A). When compared with human hippocampus, 91% of the expressed miRNAs in our primary mouse neurons were also detected in the brain tissue (Figure 1B). When we compared all three groups, 119 miRNAs were commonly detected, including various neuronal expressed miRNAs (Figure 1C) such as miR-128-3p, miR-132, miR-134-5p, miR-135a-5p, miR-181a/c, miR-212-3p and miR-218-5p. Of note, only 8 miRNAs were exclusively detected in primary hippocampal neurons (Figure 1C; miR-44b-5p, miR-422a, miR-551b-5p, miR-601, miR-659-3p, miR-661, miR-672-5p and miR-1244). Then, we focused on the cell expression of the 160 detected miRNAs in primary neurons, 158 miRNAs were identified as highly enriched in neurons, including miR-128, miR-132, miR-134 and miR-218 [9] (Figure 1D, top panel). A higher proportion of non-neuronal miRNA was detected in hippocampal tissue from mice (Figure 1D, middle panel) and human brain tissue (Figure 1D, bottom panel), 3% in both data sets.

**Table 1.** Characteristics of primary hippocampal neurons and main cell lines used on current manuscript. In current study, 1e6 hippocampal neurons were cultured for 10 Days. Immortalized cell lines were used 2 days after plated, and at a maximum confluence of 80%.

|  | Primary<br>hippocampal<br>neurons | HT22 | N2A | SHSY5Y |
| --- | --- | --- | --- | --- |
| *Species* | Mus musculus-C57BL6 | Mus musculus | Mus musculus-Albino Strain A | Homo sapiens-4 years old<br>female |
| *Cell origin* | Embryonic Day<br>18- Hippocampus | HT4 derived cell-Immortalized with<br>SV 40T virus<br>(hippocampus) | Spontaneous<br>tumour | Bone marrow<br>(patient with a<br>diagnosis of<br>neuroblastoma) |
| *Immortalized* | No | Yes | Yes | Yes |
| *Main*<br>*Applications* | Ex vivo neuronal<br>function | Glutamatergic<br>neurotoxicity | Neuronal<br>differentiation | Adrenergic cells. |

**Figure 1.**
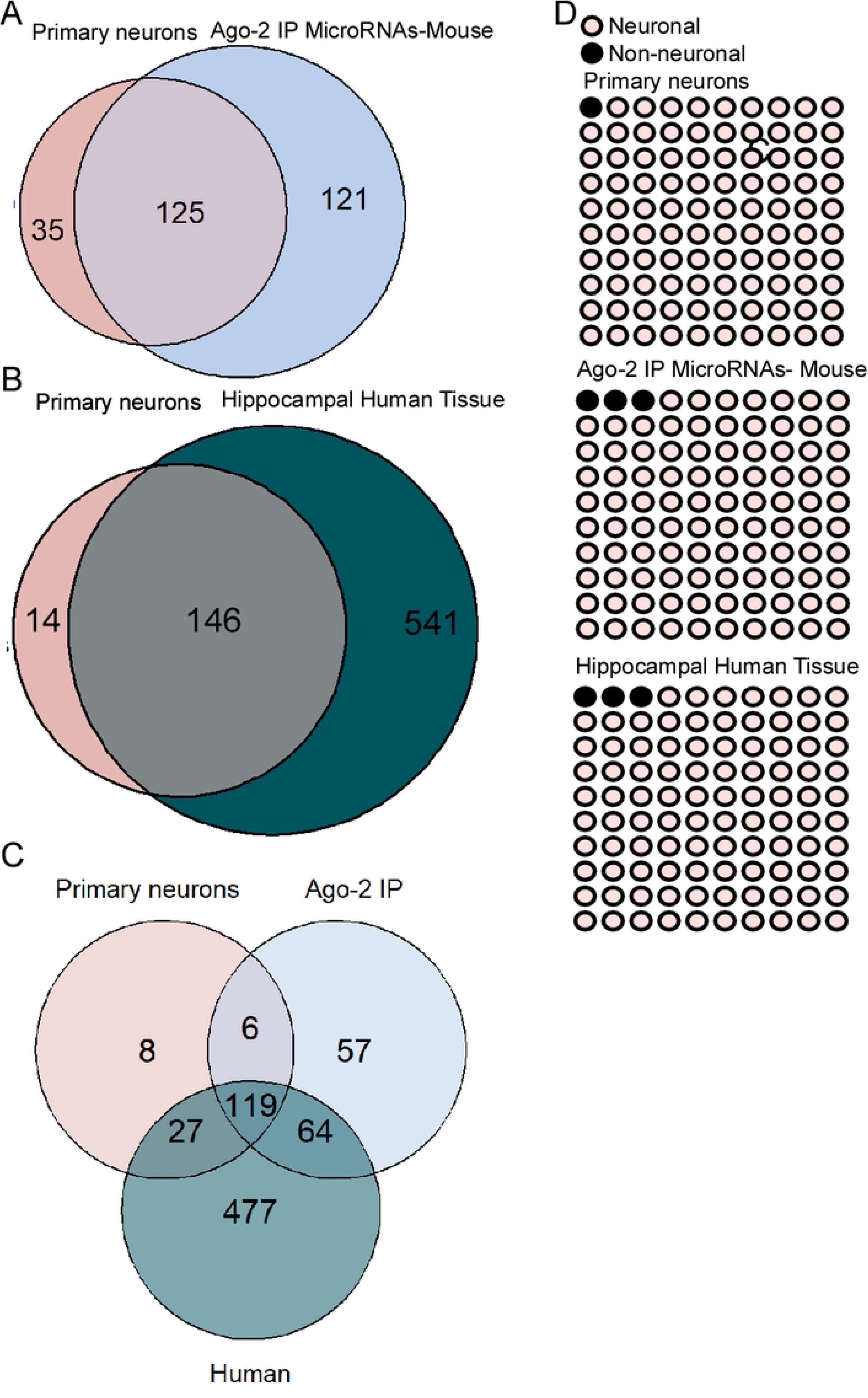
Primary hippocampal neurons have a similar microRNA profile than adult hippocampus. A,B,C) Venn diagrams show the overlap on the number of microRNAs between mouse hippocampal primary neurons, and Ago-2 bound microRNAs from the adult mouse hippocampus (A), between human control hippocampal neurons (B) and between the three tisuue types (C). D) Dot plots (10 x 10) show the proportion of neuronal enriched microRNAs in primary hypocampal neurons (top), hippocampi from adult mouse (middle), and human hippocampus (bottom).

### miRNA profiles in primary hippocampal neurons and HT22, N2A, and SH-SY5Y cell lines

Next, we evaluated the extent to which the three immortalised cell lines overlapped with the miRNA expression profile in our primary hippocampal neurons. The miRNA profiling was carried out using the qPCR-based Open Array Platform (Figure 2A). Principal Component Analysis (PCA) using the total miRNA datasets from each cell type showed separation of primary hippocampal neurons, HT22, N2A, and SH-SY5Y into four distinctive groups (Figure 2B). We detected 310 miRNAs in the four cell types (Figure 2C). Of these, 32% (98 miRNAs) were commonly detected in all the cell types. Each cell type expressed a unique subset of miRNAs; 13 miRNAs were detected only in the primary hippocampal neurons, 7 in HT22, 20 in N2A, and 61 in SH-SY5Y (Figure 2C). Then, we compared the primary hippocampal neurons with each cell line individually, the SH-SY5Y and N2A cell lines displayed the highest overlap with primary neurons, with 10 and 9 miRNAs respectively. HT22 and the primary hippocampal neuron had only 1 miRNA exclusively detected in both cell types (Figure 2C).

**Figure 2.**
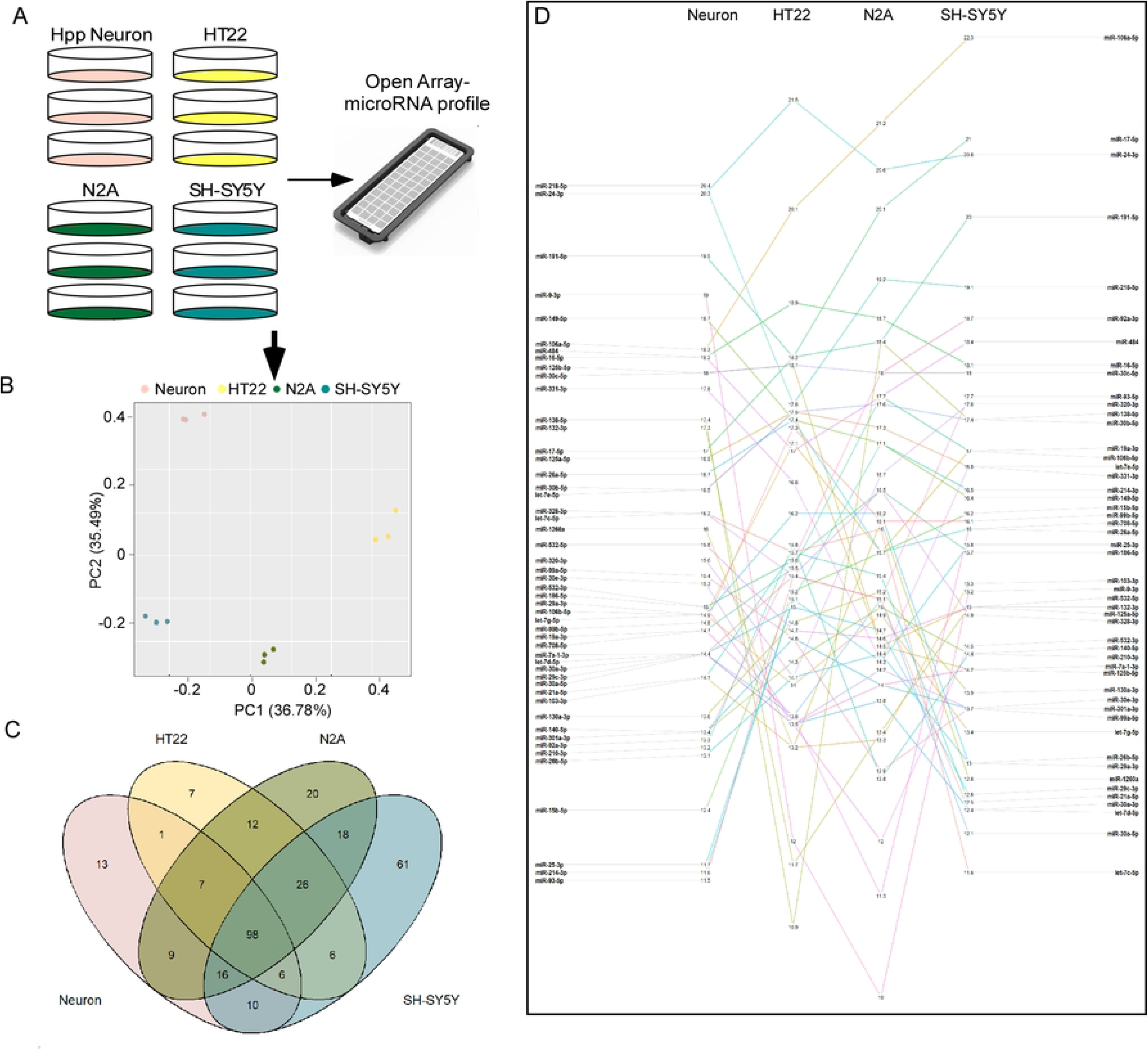
MicroRNA profile in primary hippocampal neurons, HT22, N2A and SH-SY5Y cell lines. A,B) Schematic of the procedures carried out in carrying study (A), and Principal Component Analysis (PCA,B) represent the clusters of the four cell lines. C) Volcano Plot represents overlap between the four cell lines. D) Slope graph illustrating the relative expression of the 50 most abundant microRNAs present in all four cell types. Relative expression is defined as 35 - mean Ct.

When we evaluated the level of miRNA expression in all cell types (Figure 2D), miRNAs such as miR-16-5p, miR-30c-5p, miR-125a-5p have similar expression levels in the four cell types (Figure 2D). In contrast, miR-9-3p, miR-132-3p, miR-138-5p miR-218-5p were highly expressed in neurons compared to the immortalized cell lines (Figure 2D).

### Neuron-enriched miRNAs in primary neurons and cell lines

Next, we explored the identities and relative expression of the 98 commonly detected miRNAs across the cell types (Figure 3A,B). Primary hippocampal neurons showed the highest expression of known neuron-enriched miRNAs compared to the immortalized cell lines, including miR-128, miR-132, miR-212 and miR-218 (Figure 3B). In contrast, primary neurons displayed low levels of miRNAs involved in cell proliferation, including the miR-17/92 cluster (miR-27, miR-18 and miR-92) [47] (Figure 2B). The primary neurons also displayed high expression of miRNAs linked to neurological disorders such as epilepsy, including miR-129a, miR-134, miR-135a and miR-146a [30]. In contrast, miR-129 was not detected in HT22 or N2A, miR-134 was not detected in HT22, miR-135a was not detected in N2A, and miR-146a had the lowest expression in HT22 or N2A (Figure 3B).

**Figure 3.**
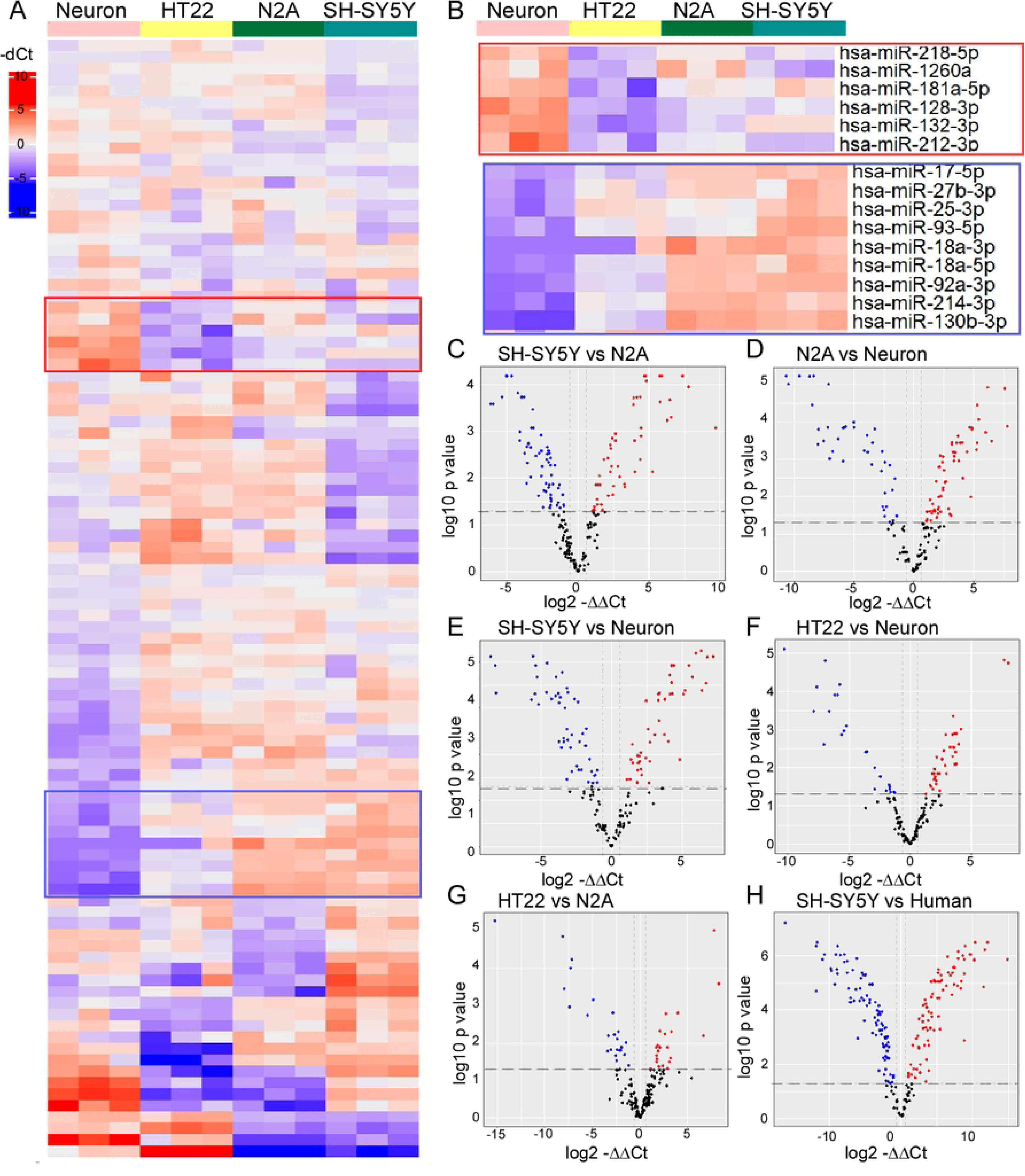
Comparison of gene expression between the cell lines. A) Heat map of the 98 microRNAs expressed in all the four cell types, each line represents an independent sample. Red: high expressed, purple: low expression. B) Amplified Heat map of the selected microRNAs highly expressed in primary neurons (top) or low expressed (bottom). C-H) Volcano plot shows the log2 FC (Fold changes) indicated by the mean expression level for each gene and each dot represents one gene. C) SH-SY5Y versus N2A, D) N2A versus hippocampal neurons, E) SH-SY5Y versus hippocampal neurons, F) HT22 versus hippocampal neurons, G) HT22 versus N2A and H) SH-SY5Y versus human brain sample.

### Differential expression of miRNAs in cell lines compared to primary hippocampal neurons

To further characterize the basal miRNA profile between the four cell types we performed differential gene expression analyses. Comparing first the two neuroblast cell lines (N2A and SH-SY5Y), we found 200 miRNAs detected with 59 miRNAs with lower expression and 50 miRNAs with higher expression levels in N2A compared to SH-SY5Y (Figure 3C). The number of commonly detected miRNAs between primary hippocampal neurons and both neuroblast cell lines was similar (163 and 166 miRNAs in N2A and SH-SY5Y respectively; Figure 3 D,E). We detected 142 miRNAs commonly detected between HT22 and primary hippocampal neurons (Figure 3F). From these commonly detected miRNAs, HT22 showed the highest level of similarly-expressed miRNAs, with 57% of the miRNAs expressed at a similar level, followed by SH-SY5Y cells, with 43% of miRNAs expressed at a similar level, and 38% of miRNAs expressed at a similar level between N2A and primary hippocampal neurons (Figure 3D,E,F).

To further understand if these similarities are driven by species or cell type, we compared both murine immortalized cell lines, HT22 and N2A (Figure 3G) and the human cell line (SH-SY5Y) to previously published human data [48] (Figure 3H). This revealed 72% of the detected miRNAs were commonly expressed between HT22 and N2A immortalized cell lines (Figure 3G) while 86% of miRNAs were commonly detected between the SH-SY5Y cell line and human tissue (Figure 3H).

### High correlation between miRNA profile in HT22 cells and primary mouse hippocampal neurons

To extend these insights, we evaluated the correlation of the full miRNA profiles between primary neuronal cell cultures and the three immortalized cell lines. This determined that HT22 cells had the highest correlation level (r = 0.735, Figure 4A), while N2A had the lowest correlation level (r = 0.564 Figure 4B), with the SH-SY5Y cell profile falling in an intermediate range (r = 0.715. Figure 4C). We further explored some of the contributors to these correlations (Figure 4). When comparing HT22 cells to primary neuronal cells, miRNAs involved in neuronal function (e.g. miR-9, miR-128, miR-132, miR-135a, miR-135b) and inflammation (miR-146a, miR-212) had lower expression levels (Figure 4A), while miRNAs involved in development and cell division (for example miR-18, miR-27a/b, miR-31, miR-106a/b and miR-222) had higher expression levels in HT22 than in primary neuronal cells cultures (Figure 4A). Similar results were obtained for N2A cells with miR-128, miR-132 or miR-134 displaying lower expression levels in N2A (Figure 4B). Interestingly, let-7e, miR-151, miR-335 or miR-378, which are involved in neuronal differentiation, have higher expression levels in N2A compared to primary cell cultures (Figure 4B).

**Figure 4.**
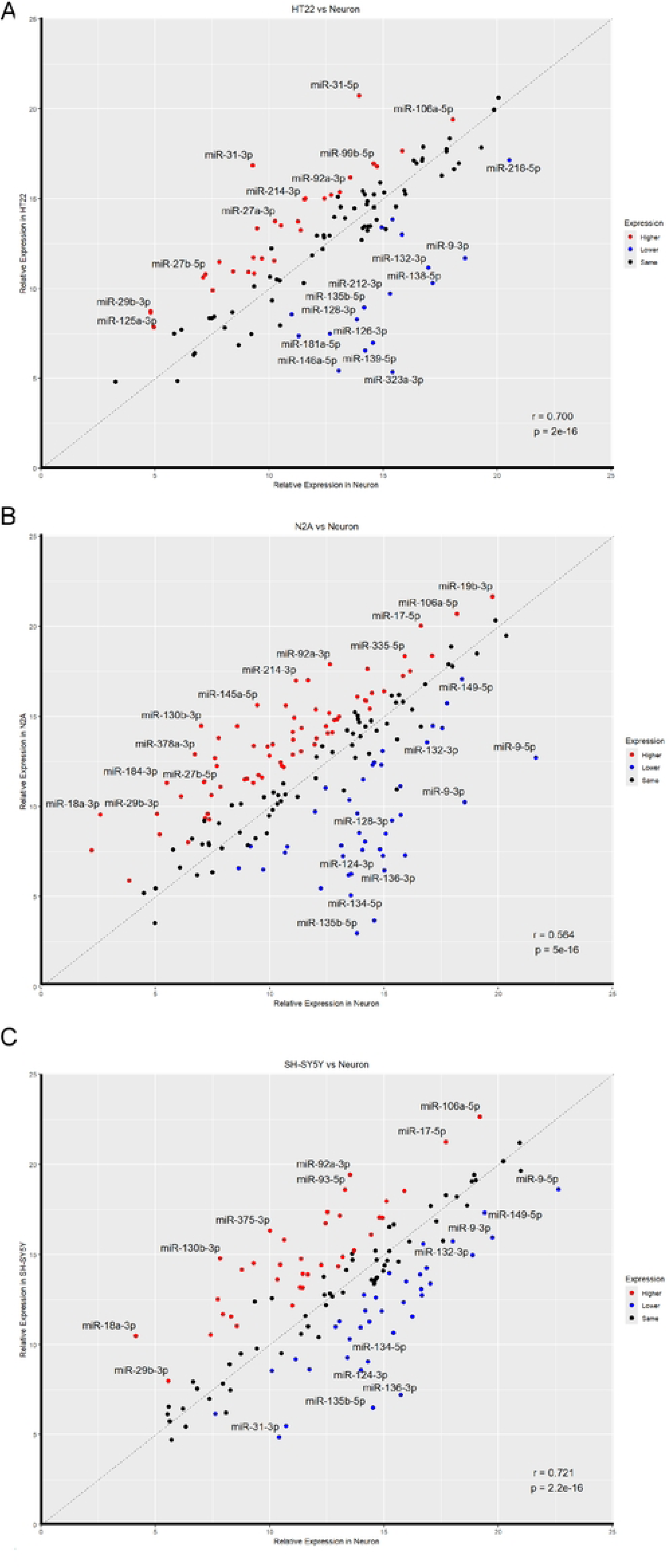
Correlations between the expression levels of the individual microRNAs between the primary hippocampal neurons and A) HT22 (r=0.735); B) N2A (r=0.564); and C) SH-SY5Y (r=0.715).

We next compared the human neuroblastoma SH-SY5Y to the primary neuronal cell line. Here we found miRNAs expressed in adult brain and involved in neuronal function such as miR-124, miR-134 or miR-135a/b also had a lower expression level in SH-SY5Y compared to the primary cell line (Figure 4C) whereas miR-27a, miR-103 or miR-140 had a higher expression level in SH-SY5Y compared to primary cell lines.

Finally, we explored the expression of additional miRNAs in each cell line. Here, we found several miRNAs have similar levels in all cells, for example, miR-30b and miR-30c (Figure 5A). MiR-125b and miR-29a have similar levels only between HT-22 and neurons (Figure 5B). While miR-106b and miR-28 have similar levels only between N2A and neurons (Figure 5C) and miR-16 and miR-29 have similar expression levels only between SH-SY5Y (Figure 5D). Together, the results represent a comprehensive mapping of the basal miRNA expression profiles in commonly used neuronal cell lines which may be useful to inform choice of cell line in miRNA research on the brain.

**Figure 5.**
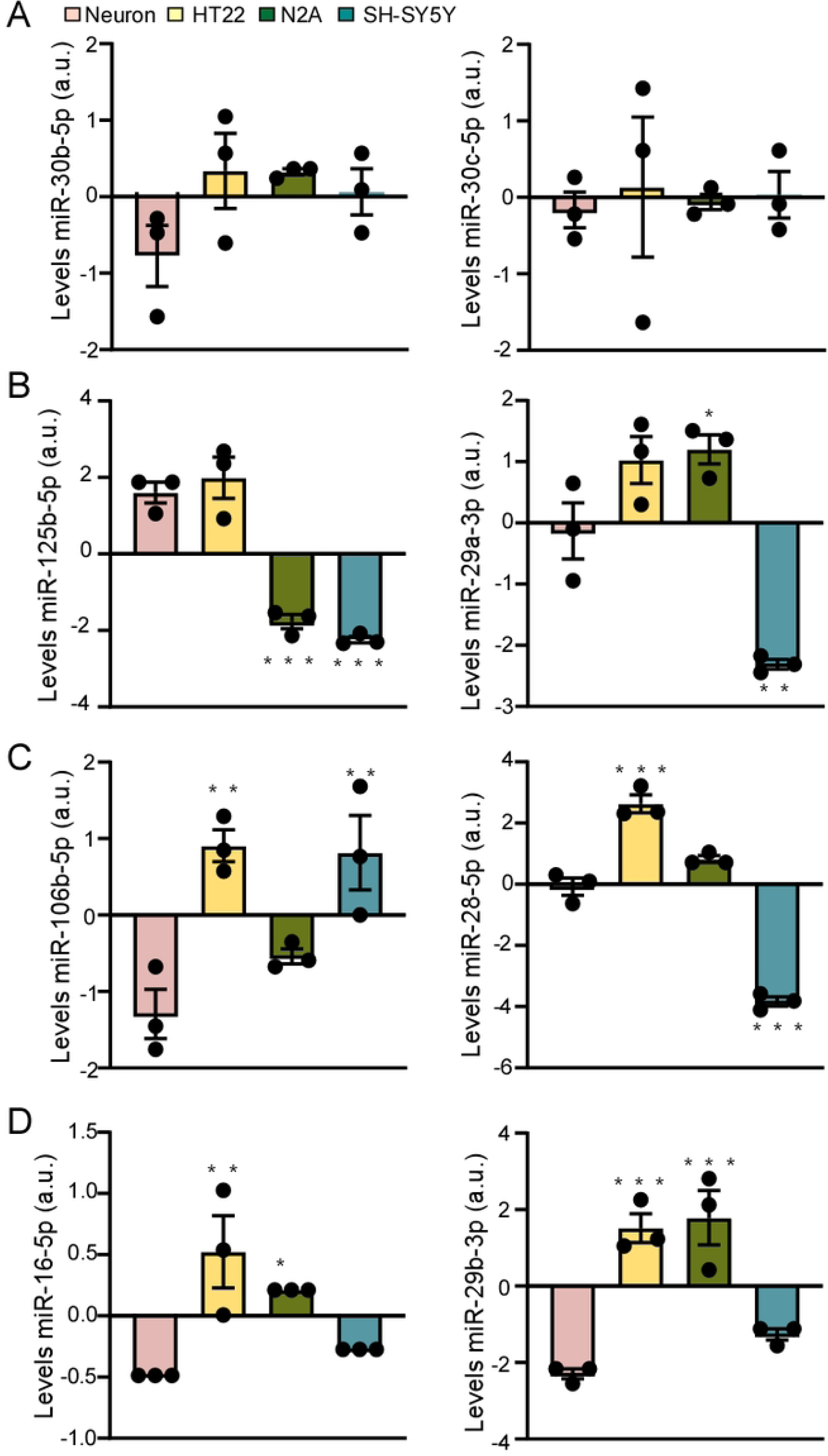
Individual microRNA levels in the four cell lines, primary hippocampal neurons (pink), HT22 (yellow), N2A (green) and SH-SY5Y (blue). A) miR-30b and miR-30c. B) miR-125b and miR-29a. C) miR-106b and miR-26. D) miR-16 and miR-29b. *p< .0.05; **p<0.01; ***p< 0.001.

## Discussion

Immortalized neuronal cell lines are invaluable tools to test the targets and function of key miRNA as well as test the properties of experimental drugs [49]. However, the suitability of cell lines for neuronal miRNA studies has often neglected an assessment of their basal miRNA expression and its relevance to primary neurons or the brain. Here, we compared three well-used immortalized neuronal cell lines, HT22, N2A and SH-SY5Y to primary mouse hippocampal neurons to evaluate their similarities and differences and support decision-making on the best cell line to study specific miRNAs or molecular mechanisms before moving to the more expensive primary cell cultures or *in vivo* experiments. Overall, the study supports the general suitability of each cell line as well as identifying profiles which may favour the selection of one cell type over another, depending on the research question. The findings provide a useful resource for experimental planning including assessing the therapeutic potential of miRNAs for neurologic diseases such as epilepsy.

A key starting point in the current study was to compare cultured primary neurons to the miRNAs that are active in the adult mouse brain. We observed that 79% of miRNAs expressed in our mouse primary hippocampal neurons were also present in the RISC complex in the adult mouse hippocampus [38], with similar results found when compared with human tissue [46]. Thus, cultured mouse neurons are a suitable proxy for the adult brain providing a benchmark for the later comparison to the cell line profiles. The small discrepancies between primary neurons and the brain (hippocampus) may reflect maturity differences or suggest a pool of miRNAs is not Ago-2 bound in the cultured neurons, perhaps as they require regulatory signals such as neuronal activity [50]. Differences may also be due to the influence or abundance of glial cells, which may be lower in the primary hippocampal neurons. Our mouse primary hippocampal cell cultures have a high (approximately 80%) shared profile of miRNAs with results from rat primary hippocampal cells [51]. This includes similar expression of miR-125a, miR-125b, miR-184, miR-195, miR-214 and miR-384, which in rat and mouse hippocampal neurons are in the highest expression group.

Furthermore, miR-22, miR-24, miR-26, miR-29a, miR-124 and miR-129, are within the lowest expressed miRNAs; and miR-103, miR-130a, miR-130b, miR-134, miR-376 and miR-449 are in the group with intermediate expression [44]. However, this differed from mouse cortical primary neurons, where the highest expressed miRNAs are miR-124, miR-132 and miR-135b [9], suggesting that the brain region may influence miRNA levels. Importantly, a larger number of miRNAs were only detected in the human tissue. This may reflect, however, that RNAseq was used to generate those data which allows for a broader range of detection than the Open Array Platform.

While cell lines cannot fully replicate the characteristics of primary hippocampal neurons, the present study reveals that HT22, N2A and SH-SY5Y cells express miRNAs that make them suitable for the analysis of neuronal and/or brain-related processes. Specifically, each cell line expressed a large number of the same miRNAs as are present in mouse primary hippocampal neurons. For example, N2A cells have similar miR-106b and miR-28 expression levels than primary hippocampal neurons, and SH-SY5Y cells have similar miR-16 and miR-29 expression levels than primary hippocampal neurons. Similar to both neuroblastoma cell lines, miR-125b and miR-29a have similar expression levels between HT22 and primary hippocampal neurons. Since both miRNAs are involved in regulating processes of neuronal death [52–55], HT22 cells may be suitable for analysing neuronal apoptosis regulated by miRNAs.

A key observation in the present study was that while miRNA profiles show substantial overlap with primary mouse neurons, they tend to express lower levels of adult-enriched miRNAs. This is not surprising and suggests researchers should be wary that the cell lines are a better model for neurodevelopment and may retain tumor-related miRNA profiles. We note, however, that the cells were not differentiated and it would be interesting to monitor how miRNA profiles develop upon differentiation to a more mature neuron-like state. The composition of miRNAs for each cell line is not fixed, and could be adjusted experimentally, for example by differentiation or introduction or deletion of miRNAs or miRNA maturation components. It would be interesting to observe if such manipulations (e.g. of miR-124, miR-134) resulted in acquisition of neuronal projection features such as axons or dendrites.

Finally, our study did not find a large effect of species, with most miRNA profiles quite conserved between human and mouse lines. This supports the relevance of the mouse lines for miRNA functions in human diseases. There are, however, human-specific miRNAs, many of which are expressed uniquely in the brain [56], which should be considered according to the research question. Regardless, these findings suggest that the cellular function of interest should be the primary decision point in choosing the cellular line rather than the species origin.

There are some limitations in the present study. First, we used a qPCR-based system to profile miRNAs. The OpenArray platform has a pre-selected number of miRNAs such that we have not captured the full repertoire of miRNAs. We will also have missed isomiRs and editing, which are important for miRNA function [57]. Second, we did not include human primary (e.g. iPSC) neurons which would provide a useful additional comparator. Third, our study only evaluated the basal levels of miRNA expression and cannot be used to infer how the cell line miRNA profiles may respond to stressors.

In conclusion, we used a low-input profiling platform to screen miRNAs in three common cell lines and compared findings to primary neurons. We found overall a high degree of correlation indicating these lines are suitable ‘models’ for the identification of miRNA-related processes and their target genes. However, specific miRNA levels varied between lines with some miRNA absent while each line varied in terms of aligning to primary neurons. The results provide a catalogue of miRNA expression which researchers can use to select the most appropriate cell type for basic and applied miRNA research, including for studies on epilepsy.

## Authors contribution

RM carried out analysis, JVS carried out analysis original RNA sequencing, ASR and EJM carried out cell cultures and Open Array, CM and TE contributed to the work on Ago-2 bound miRNA, MV supported analysis of Ago RNA sequencing. DCH and EJM conceived the study and co-wrote the manuscript.

## Acknowledgements

We would like to thank Carsten Culmsee (Philipps-Universität Marburg) for the HT22 cell line and Jochen Prehn (Royal College of Surgeons in Ireland) for the N2A and SH-SY5Y cell lines. We also thank James Mills for help with analysis of the miRNA data from human brain tissue. This publication has emanated from research conducted with the financial support of the European Union’s ‘Seventh Framework’ Programme (FP7) under Grant Agreement no. 602130. Additional support is from Health Research Board under Grant number ILP-POR-2022-029, Wellcome Trust-IISF call under Grant number Ph2IISF-16585, and Taighde Éireann – Research Ireland, under Grant number 17/CDA/4708, 22/FFP-P/11333 and 21/RC/10294_P2 at FutureNeuro Research Ireland Centre for Translational Brain Science.

